# Epitope classification and RBD binding properties of neutralizing antibodies against SARS-CoV-2 variants of concern

**DOI:** 10.1101/2021.04.13.439681

**Authors:** Ashlesha Deshpande, Bethany D. Harris, Luis Martinez-Sobrido, James J. Kobie, Mark R. Walter

## Abstract

Severe acute respiratory syndrome coronavirus-2 (SAR-CoV-2) causes coronavirus disease 2019 (COVID19) that is responsible for short and long-term disease, as well as death, in susceptible hosts. The receptor binding domain (RBD) of the SARS-CoV-2 Spike (S) protein binds to cell surface angiotensin converting enzyme type-II (ACE2) to initiate viral attachment and ultimately viral pathogenesis. The SARS-CoV-2 S RBD is a major target of neutralizing antibodies (NAbs) that block RBD - ACE2 interactions. In this report, NAb-RBD binding epitopes in the protein databank were classified as C1, C1D, C2, C3, or C4, using a RBD binding profile (BP), based on NAb-specific RBD buried surface area and used to predict the binding epitopes of a series of uncharacterized NAbs. Naturally occurring SARS-CoV-2 RBD sequence variation was also quantified to predict NAb binding sensitivities to the RBD-variants. NAb and ACE2 binding studies confirmed the NAb classifications and determined whether the RBD variants enhanced ACE2 binding to promote viral infectivity, and/or disrupted NAb binding to evade the host immune response. Of 9 single RBD mutants evaluated, K417T, E484K, and N501Y disrupted binding of 65% of the NAbs evaluated, consistent with the assignment of the SARS-CoV-2 P.1 Japan/Brazil strain as a variant of concern (VoC). RBD variants E484K and N501Y exhibited ACE2 binding equivalent to a Wuhan-1 reference SARS-CoV-2 RBD. While slightly less disruptive to NAb binding, L452R enhanced ACE2 binding affinity. Thus, the L452R mutant, associated with the SARS-CoV-2 California VoC (B.1.427/B.1.429-California), has evolved to enhance ACE2 binding, while simultaneously disrupting C1 and C2 NAb classes. The analysis also identified a non-overlapping antibody pair (1213H7 and 1215D1) that bound to all SARS-CoV-2 RBD variants evaluated, representing an excellent therapeutic option for treatment of SARS-CoV-2 WT and VoC strains.

## Introduction

A robust humoral immune response to severe acute respiratory syndrome coronavirus-2 (SARS-CoV-2) infection is essential to control viral infection (1,2). Human neutralizing antibodies (NAbs) that bind to the receptor binding domain (RBD) of the SARS-CoV-2 Spike (S) protein and block ACE2 binding have shown efficacy as therapeutics and are also essential for vaccine efficacy (3–6). The SARS-CoV-2 S protein is a trimer, where the protomers undergo conformational changes essential for viral infectivity (7,8). Specifically, the RBD moves from a “down” conformation, where the ACE2 binding site is buried in the S trimer, to an “up” conformation where the ACE2 binding site is solvent accessible and competent for cell surface ACE2 binding (**Fig. 1**). Presumably, the “down” conformation of RBD “hides” the ACE2 binding site in the closed trimeric arrangement to, at least partially, protect the RBD, which is essential for virus cell attachment, from the host’s immune response.

**Fig. 1.**
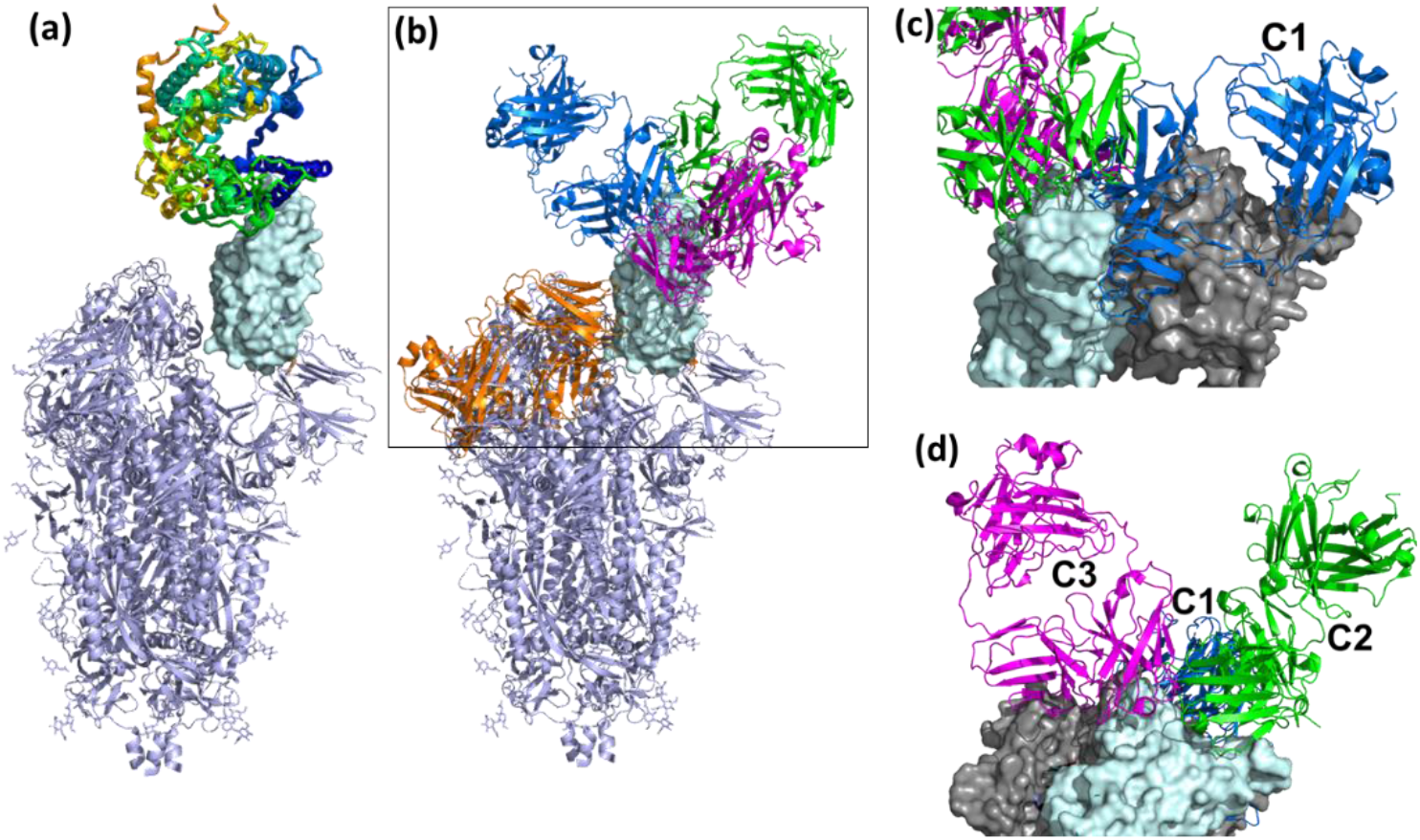
Overall view of ACE2 and NAbs binding to SARS-CoV-2 S protein. **(a)** Ribbon diagram of the SARS-CoV-2 S protein, with one RBD in the up conformation (cyan surface) interacting with ACE2. **(b)** General location of NAb binding epitopes, where an example C1 NAb is colored blue, C2 green, C3 magenta and C4 orange. The figure highlights that extensive conformational changes are required before C4 NAbs can bind to S. (**c**) View of two RBDs from an S trimer (cyan and grey surfaces) in the down conformation, to show that C1 NAbs cannot bind to their epitope while the RBD is in the down conformation, due to steric hinderance. (**d**) Alternative view of adjacent down RBDs in the S trimer showing C2 and C3 NAb epitopes are accessible while RBD is in the down conformation and that some NAbs can bind across an RBD-RBD trimer interface and interfere with RBD opening to bind ACE2.

Three-dimensional structural studies have played a large role in our understanding of SARS-CoV-2 S RBD-specific NAb epitopes (**Table 1**). To date, NAbs targeting SARS-CoV-2 RBD have been classified into four structural groups, C1-C4 (**Fig. 1**), based on their ability to bind to up or down conformations of the SARS-CoV-2 RBD, and by their overlap with the ACE2 binding site (9). The C1 class is defined by NAbs that bind within the RBD ACE2 binding site, essentially mimicking ACE2 binding, and thus only bind to RBD when it is in the up conformation. The C1 class contains conserved ACE2-blocking NAbs, often composed of heavy chains encoded by the VH3-52 and VH3-66 gene segments (10). C2 NAbs block the ACE2 binding site yet can bind to RBD in its up and down conformations. C3 NAbs bind largely outside the ACE2 binding site of RBD in both up and down conformations, while a fourth class (C4) binds further from the ACE2 binding site and only the epitope is only accessible once SARS-CoV-2 S undergoes large conformational changes (11). Overall, a variety of NAb/RBD structures have been completed and their interactions defined (see refs. in Table 1).

| CLASS 1: (n=17) |  |  |  |  |  |  |  |  |  |  |  |
| --- | --- | --- | --- | --- | --- | --- | --- | --- | --- | --- | --- |
| KD (pM)* | IC50 (ng/mL)* | # | pdb | name | unknown | HC BSA | LC BSA | HC of Tot. | HC+LC | M. | Ref. |
|  |  | 1 | 6xc2 | cc12.1 |  | 766.7 | 574.9 | 0.57 | 1341.6 | X | (33) |
| 186 | 120 / 170 |  |  |  | 1212F2 |  |  |  |  |  |  |
|  |  | 2 | 6xc4 | cc12.3 |  | 676.9 | 186.2 | 0.78 | 863.1 | X | (33) |
|  |  | 3 | 6xe1 | CV30 |  | 713.5 | 271.3 | 0.72 | 984.8 | X | (34) |
|  |  | 4 | 7bz5 | B38 |  | 691.8 | 501.2 | 0.58 | 1193 | X | (35) |
| 11,600 | 323 / 204 | 5 | 7c01 | CB6 |  | 736.1 | 312.6 | 0.70 | 1048.7 | X | (30) |
| 738 | 768 / 107 |  |  |  | 1214F4 |  |  |  |  |  |  |
| 122 | 210 / 290 |  |  |  | 1212D5 |  |  |  |  |  |  |
|  |  | 6 | 7cdi | P2C-1F11 |  | 726.5 | 229.3 | 0.76 | 955.8 | X | (36) |
| 18 | 1127 / 168 |  |  |  | 1214E9 |  |  |  |  |  |  |
| 13,500 | 460 / 140 |  |  |  | 1212D4 |  |  |  |  |  |  |
|  |  | 7 | 7cjf | - |  | 771 | 430.8 | 0.64 | 1201.8 | X | - |
|  |  | 8 | 7jmo | COVA2-04 |  | 757 | 369.4 | 0.67 | 1126.4 | X | (37) |
|  |  | 9 | 7k8m | C102 |  | 802.4 | 231 | 0.78 | 1033.4 | X | (9) |
|  |  | 10 | 7kfv | C1A-B12 |  | 809.1 | 375.8 | 0.68 | 1184.9 | X | (38) |
|  |  | 11 | 7kfw | C1A-B3 |  | 776.6 | 371.2 | 0.68 | 1147.8 | X | (38) |
|  |  | 12 | 7kfx | C1A-C2 |  | 854.9 | 353.3 | 0.71 | 1208.2 | X | (38) |
| 121 | 490 / 380 |  |  |  | 1212D6 |  |  |  |  |  |  |
| 7.2 | 900/100 |  |  |  | 1206D12 |  |  |  |  |  |  |
| 161 | 240 / 550 |  |  |  | 1213F2 |  |  |  |  |  |  |
|  |  | 13 | 7kfy | C1A-F10 |  | 797.8 | 367.8 | 0.68 | 1165.6 | X | (38) |
|  |  | 14 | 7chf | BD-604 |  | 712.6 | 397.2 | 0.64 | 1109.8 | X | (39) |
| 2,470 | 872 / 112 |  |  |  | 1212C8 |  |  |  |  |  |  |
|  |  | 15 | 7ch5 | BD-629 |  | 818.2 | 193.3 | 0.81 | 1011.5 | X | (39) |
|  |  | 16 | 7chb | BD-236 |  | 656.3 | 425 | 0.61 | 1081.3 | X | (39) |
|  |  | 17 | 6xcm | C105 |  | 635.2 | 258.1 | 0.71 | 893.3 | EM | (9) |
| CLASS-1 Distinct (n=5) |  |  |  |  |  |  |  |  |  |  |  |
| KD (pM)* | IC50 (ng/mL)* | # | pdb | name | unknown | HC BSA | LC BSA | HC of Tot. | HC+LC | M | Ref. |
|  |  | 18 | 6xkq | CV07-250 |  | 422 | 544.8 | 0.44 | 966.8 | X | (26) |
|  |  | 19 | 6xdg | RGN10933 |  | 760.8 | 187.9 | 0.80 | 948.7 | X | (4) |
|  |  | 20 | 7jx3 | S2H14 |  | 502.7 | 408.2 | 0.55 | 910.9 | X | (40) |
|  |  | 21 | 7k9z | 298Fab |  | 409.5 | 302.6 | 0.58 | 712.1 | X | - |
|  |  | 22 | 7jmp | COVA2-39 |  | 616.2 | 143 | 0.81 | 759.2 | X | (37) |
| 1,180 | 66 / 49 |  |  |  | 1213H7 |  |  |  |  |  |  |
| CLASS 2: (n=13) |  |  |  |  |  |  |  |  |  |  |  |
| KD (pM)* | IC50 (ng/mL) | # | pdb | Name | unknown | HC BSA | LC BSA | HC of Tot. | HC+LC |  |  |
|  |  | 23 | 7jv2 | S2H13 |  | - | - | - | - | EM |  |
|  |  | 24 | 7k8u | C104 |  | - | - | - | - | EM | (9) |
| 21.9 | 117 / 85 |  |  |  | 1218A6 |  |  |  |  |  |  |
|  |  | 25 | 7k8y | C121 |  | 741.9 | 56.4 | 0.93 | 798.30 | EM | (9) |
| 9,630 | 777 / 1,312 |  |  |  | 1206D1 |  |  |  |  |  |  |
| 104 | 10 / 22- |  |  |  | 1212C2 |  |  |  |  |  |  |
| 841 | 31 / 26 |  |  |  | 1212F5 |  |  |  |  |  |  |
| 3,530 | 300 / 1,380 |  |  |  | 1207F10 |  |  |  |  |  |  |
|  |  | 26 | 7k8s | C002 |  | 645.2 | 181.9 | 0.78 | 827.10 | EM | (9) |
|  |  | 27 | 7k90 | C144 |  |  |  |  |  | EM | (9) |
|  |  |  |  | C144A |  | 684.9 | 102.4 | 0.87 | 787.30 |  |  |
|  |  |  |  | C144B |  | 693 | 103.1 | 0.87 | 796.10 |  |  |
|  |  |  |  | C144AB |  | 1088 | 118.7 | 0.90 | 1206.70 |  |  |
|  |  |  |  | C144C |  | 698.4 | 100.2 | 0.87 | 798.60 |  |  |
|  |  | 28 | 6xey | Ab2-4 |  | 678.3 | 160.8 | 0.81 | 839.10 | EM | (27) |
|  |  | 29 | 7che | C368-2 |  | 600.2 | 142.5 | 0.81 | 742.70 | X | (39) |
| 77 | 226 / 59 |  |  |  | 1215D1 |  |  |  |  |  |  |
| 1,400 | 210 / 310 |  |  |  | 1207B4 |  |  |  |  |  |  |
| 658 | 650 / 180 |  |  |  | 1206A5 |  |  |  |  |  |  |
|  |  | 30 | 7bwj | P2B-2F6 |  | 481.5 | 142.6 | 0.77 | 624.10 | X | (36) |
|  |  | 31 | 7k8w | C119 |  |  |  |  |  | EM | (9) |
|  |  |  |  | C119A |  | 613.8 | 346.8 | 0.64 | 960.60 |  |  |
|  |  |  |  | C119B |  | 617.3 | 271.5 | 0.69 | 888.80 |  |  |
|  |  | 32 | 7byr | Ab23 |  | 730.5 | 34.3 | 0.96 | 764.80 | EM | (41) |
| 2,470 | 872 / 112 |  |  |  | 1218C8 |  |  |  |  |  |  |
|  |  | 33 | 6xkp | CV07-270 |  | 693.5 | 106.8 | 0.87 | 800.30 | X | (26) |
|  |  | 34 | 7cdj | P2C-1A3 |  | 587.5 | 300.2 | 0.66 | 887.70 | X | (36) |
|  |  | 35 | 7k9z | 52Fab |  |  |  |  |  | X | - |
| CLASS-3: (n=4) |  |  |  |  |  |  |  |  |  |  |  |
| KD (pM)* | IC50 (ng/mL)* | # | pdb | common | unknown | HC BSA | LC BSA | HC of Tot. | HC+LC |  |  |
|  |  | 36 | 7k8v | C110A |  | 361 | 268 | - | - | EM | (9) |
|  |  |  |  | C110B |  | 249 | 320 | - | - | EM | (9) |
|  |  | 37 | 7k8z | C135A |  | 273 | 135 | - | - | EM | (9) |
|  |  |  |  | C135B |  | 354 | 225 | - | - | EM | (9) |
| 8,150 | 53 / 111 | 38 | 6xdg | RGN10987 |  | 512.9 | 112.7 | 0.82 | 625.6 | X | (4) |
|  |  | 39 | 7jx3 | S309 |  | 608.2 | 167.5 | 0.78 | 775.7 | X | (42) |
| CLASS-4 (n=6) |  |  |  |  |  |  |  |  |  |  |  |
| KD (pM)* | IC50 (ng/mL)* | # | pdb | common | unknown | HC BSA | LC BSA | HC of Tot. | HC+LC |  |  |
|  |  | 40 | 6w41 | CR3022 |  | 593.4 | 397.3 | 0.60 | 990.7 | X | (10) |
|  |  | 41 | 6zer | EY6A |  | 573 | 373.4 | 0.61 | 946.4 | X | (43) |
|  |  | 42 | 7cah | H014 |  | 673.5 | 287.6 | 0.70 | 961.1 | EM | (28) |
|  |  | 43 | 7jva | S2A4 |  | 375.5 | 431.2 | 0.47 | 806.7 | EM | (40) |
|  |  | 44 | 7jmw | COVA1-16 |  | 625.3 | 153.4 | 0.80 | 778.7 | X | (29) |
|  |  | 45 | 7jx3 | S304 |  | 491.3 | 368.7 | 0.57 | 860 | X | (40) |
| 500 | >10,000 |  |  |  | 1215B11 |  |  |  |  |  |  |

\* values taken from reference (16)

Structural correlates of SARS-CoV-2 neutralization can inform NAb mechanisms of action and their potential susceptibility to S protein RBD variants, either alone or in combinations, both of which are observed in VoCs (12–15). Thus, there remains a great need to characterize the molecular details of NAb responses induced upon natural infection and/or by vaccination, at the patient and population levels. This is a large endeavor that ultimately will require protein modeling, combined with synergistic experimental studies and additional 3D structure determinations. As a first step towards this goal, we sought to understand the RBD binding epitopes of a panel of SARS-CoV-2 NAbs, isolated from a single convalescent patient (16). To accomplish this goal, NAb-RBD structures (n=45, Table 1) in the protein databank (pdb) were grouped into five distinct classes based on their RBD binding profiles. The 20 unknown NAbs were assigned to a NAb/RBD complex and validated using RBD variant binding and epitope mapping studies. RBD variants were chosen based on structural considerations and RBD sequence variation observed in the GISAID (17). Of the nine RBD variants tested, N501Y, E484K, and K417T disrupted binding to 65% of the NAbs evaluated. Furthermore, N501Y and E484K RBD variants exhibited essentially wild type (WT) affinity for ACE2, where WT is defined as the Wuhan-1 RBD sequence. Another RBD variant, L452R, disrupted fewer NAbs, but exhibited 2-fold higher affinity for ACE2, suggesting this mutation may enhance viral fitness through increased affinity for ACE2, as well as disrupting the host NAb response. The analysis identified a pair of potent NAbs with non-overlapping binding epitopes, 1213H7 (NT_50_ = 49 ng/mL) and 1215D1 (NT_50_ = 59 ng/mL), that bound to all nine RBD variants tested. Furthermore, 1215D1 has a novel neutralization mechanism, as it can bind to a preformed RBD-ACE2 complex, potentially allowing it to disrupt the interaction between the virus and ACE2 even after it is formed.

## Materials and Methods

### NAb-RBD complex structure classifications and assignments

Binary RBD-NAb complex structures (human NAb only), solved by X-ray crystallography and cryo-electron microscopy, were extracted from the pdb (n=45, 12-26-21). RBD surface area buried upon NAb-RBD complex formation was quantified using PISA (18) to define a RBD residue-specific NAb binding profile (BP) for each structure. Buried surface area was not calculated for some complexes (e.g. C104, pdbid = 7k8u), due to almost complete lack of sidechains for the NAbs. The NAb BPs were subjected to hierarchical clustering and a correlation-based similarity matrix was calculated using the program Morpheus (https://software.broadinstitute.org/morpheus). To evaluate the classifications, RBDs from each NAb-RBD complex were superimposed and the structures of the resulting NAb classifications compared using pymol (19). Unknown NAbs were assigned to the classified NAbs by heavy chain sequence alignment using clustal omega (20).

### RBD variant analysis

SARS-CoV-2 S protein sequences were downloaded from the GISAID (17) for analysis on 1-11-2021. We acknowledge all authors that submitted sequences to the GISAID, which are listed in (https://www.gisaid.org/). The RBD region, residues 343-532, of full-length SARS-CoV-2 S sequences (322,688 sequences) were evaluated for amino acid sequence variation at each RBD sequence position, relative to the Wuhan-1 sequence (pub med accession number NC045512.2) (21). The total number of amino acid changes at each position was counted and converted to a frequency by dividing by the total number of sequences analyzed.

### Protein Expression

SARS-CoV-2 NAbs were produced and purified as previously described (16). SARS-CoV-2 RBD (Wuhan-1) and RBD variants were produced in insect cells as RBD-FC fusion proteins (22). RBD-FC fusion proteins were purified from expression media by protein-A affinity chromatography, as previously described (16,22). Due to a sequencing error, RBD E484K-FC used for all binding studies was determined to be RBD E484R-FC. Due to the similarity of the amino acid chemistry between Arginine and Lysine, the RBD E484R provides a suitable surrogate for the RBD E484K variant. Soluble ACE2 and RBD-ACE2 fusion protein (RBD-ACE2FP) were expressed in insect cells as C-terminal his-tagged proteins. The proteins were purified from expression media by nickel affinity chromatography, followed by dialysis into 20mM HEPES, pH 7.4, 150mM NaCl for binding studies.

### Surface plasmon resonance (SPR) Binding Studies

SPR binding studies were performed using a Biacore T200 (GE life sciences). All studies were performed at 25°C in running buffer consisting of 10mM HEPES, pH 7.4, 150mM NaCl, 0.0075% P20, and 75ug/mL bovine serum albumin. Epitope mapping was performed by capturing RBD-FCs to CM-5 sensor chips using an anti-murine FC capture kit (Cytiva). Two NAbs (50nM concentrations) were sequentially injected (NAb1, followed by NAb2) for 90 seconds over each RBD-FC surface, followed by a 60 second dissociation phase. The flowrate for the epitope mapping studies was 30 μL/min. Binding levels (RU) of each NAb was recorded following the binding phase of each injection. At the end of each cycle, the surface was regenerated by a 3-minute injection of 10 mM glycine pH 1.7. Competition between NAb epitopes was defined relative to NAb1 (e.g. NAb2 binding (RU)/ NAb1 binding (RU)).

ACE2 binding affinity for RBD variants was performed by capturing RBD-FCs to CM-5 chips, as described in the epitope mapping studies. Soluble ACE2 was injected over the RBD surfaces at 5 concentrations (300 nM, 75 nM, 18.75 nM, 4.7nM, and 1.2 nM) at a flow rate of 40 μL/min. The resulting sensorgrams were globally fit to a 1:1 model using Biacore T-200 evaluation software version 1.0 to derive binding constants.

NAb-RBD variant interactions were characterized by injecting RBD-FC Wuhan-1 reference strain, and RBD-FC variants, over NAbs captured on CM5 chip surfaces, using a human IgG capture kit (Cytiva). RBD-FCs (50nM) were injected at a flow rate of 50μL/min over the NAb surfaces for 90 seconds, followed by a 5 min. dissociation. RBD-variant binding responses (RU) were compared to the RBD reference WT protein (e.g. RU-RBD variant / RU-RBD-WT), collected at the beginning and end of the experiment and converted to percentage of WT binding.

RBD-ACE2FP-NAb epitope analysis was performed injecting RBD-ACE2FP (20nM) over each NAb, coupled as described for the NAb-RBD variant interactions. RBD-ACE2FP was injected over the surface for 90 seconds, followed by a 5 min. dissociation. The flow rate for the experiments was 50μL / min. Buffer subtracted sensorgrams were visually evaluated and RU values at 570 seconds, post injection, recorded.

## Results

### Structure of the RBD-ACE2 binding site

The structure of the SARS-CoV-2 RBD-ACE2 complex ((23), PDBID = 6m0j) was used to define RBD residues that bind to ACE2. Residues 333-526 of the 1273 amino acid SARS-CoV-2 S protein form the RBD. The RBD consists of a scaffold, conserved among different coronaviruses, and a variable receptor binding module (RBM), which positions amino acids for contact with ACE2 (24). Three RBM regions: knob (444-449 and 498-505), base (490-494 and 492-497), and tip (473-489), contact the N-terminal α-helical region of ACE2. (**Fig. 2a**). A total of 22 RBD residues bury surface area (total = 859Å^2^) and participate in 15 hydrogen bonds with ACE2, of which 11 are formed by RBD sidechains (**Figs. 2b-2e**). RBD residues Y449, N487, and Y505 form bivalent hydrogen bonds with ACE2, accounting for 6 of 10 sidechain hydrogen bonds, while K417, Y489, Q493, N498, and T500 form the five remaining sidechain hydrogen bonds with ACE2 (**Fig. 2e**). These residues are of particular interest, since RBD amino acid variants in these positions may alter the interaction of RBD with ACE2.

**Fig. 2.**
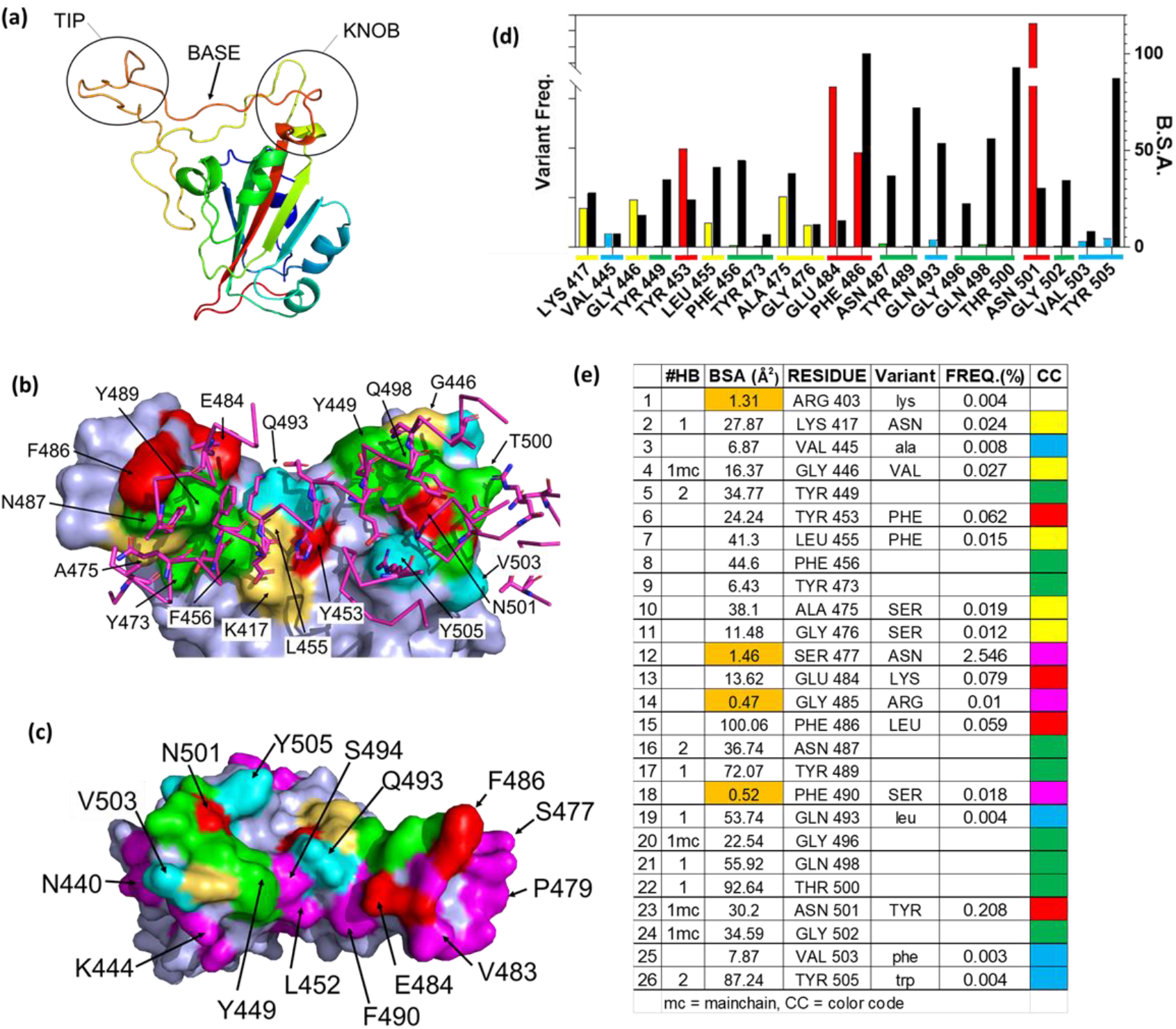
Details of the RBD – ACE2 interaction and frequency of RBD mutations. (**a**) Ribbon diagram of the SARS-CoV-2 S RBD highlighting the 3 regions (tip, base, and knob) of the RBM that bind to ACE2. (**b**) Surface representation of the RBD ACE2 binding site (pdbid 6m0j) color coded based on amino acid variability, relative to Wuhan-1 reference sequence (see **Fig. 2e** for color code). The sidechains and backbone of ACE2 amino acids that contact the RBD are shown on the figure and colored magenta. (**c**) RBD ACE2 binding surface, rotated 180 degrees relative to **Fig. 2b**, that highlights residues adjacent to the ACE2 binding site, that exhibit amino acid variation of at least 0.01% (magenta). (**d**) Graphical presentation of the frequency of RBD variant occurrences in the GISAID (left axis, colored bars), relative to buried surface area (right axis, black bars). Coloring described in Fig **2e**. (**e**) Parameters of the RBD-ACE2 binding site. All 26 RBD residues that bury >0Å^2^ of surface area in the RBD-ACE2 complex are listed along with the amount of buried surface area (BSA). Residues with BSA colored orange highlight residues that are not considered part of the ACE2 binding site. The types of hydrogen bonds made by each residue is listed under #HB, where bivalent sidechain hydrogen bonds = 2, monovalent sidechain hydrogen bonds =1, and mainchains hydrogen bonds are denoted by “mc”. Frequency that variant RBD amino acids (variant) observed in SARS-CoV-2 GISAID SARS-CoV-2 S sequence database, relative to the Wuhan-1 reference sequence. The color code (CC) for Figs. **2b**, **2c**, and **2d** is also shown.

### RBD Variant Analysis

As RBD-ACE2 interactions are essential for virus survival, we hypothesized that energetically essential ACE2 binding residues would remain invariant as the virus evolved, while residues less important for ACE2 binding, would be more susceptible to sequence variation. To test this hypothesis, the mutation frequency at each amino acid position in the RBD of SARS-CoV-2 S was compared against the Wuhan-1 reference S RBD sequence (NC045512.2). Of the 322,688 S sequences evaluated, 9 ACE2 binding residues (41%) have remained invariant (0.001% variation or less, **Fig. 2e**). These invariant residues cluster within the tip (F456, Y473, N487, Y489) and knob (Y449, G496, Q498, T500, G502) of RBD (**Figs. 2a, 2b**) where they participate in a series of contacts with ACE2 (**Figs. 2d 2e**). Two invariant residues (Y449 and N487) participate in bivalent hydrogen bonding networks with ACE2, five additional hydrogen bonds are formed with residues Y489, G496, Q498, T500, G502, while F456 and Y473 form a hydrophobic scaffold that allows ACE2 to efficiently pack against the RBD tip (**Fig. 2e**). Overall, the invariant residues account for 60% of the hydrogen bonds in the RBD-ACE2 interface. These observations suggest that ACE2 binding requirements restrain SARS-CoV-2 RBD amino acid variation at certain positions (**Figs. 2d, 2e**).

In contrast to the invariant residues, 9 ACE2 binding residues have undergone significant sequence variation (**Fig. 2**) with the largest frequencies occurring at four positions: N501Y (0.208%), E484K (0.079%), Y453F (0.062%), and F486L (0.059%). The N501Y variant, which occurs at the highest frequency, is the only residue located within the RBD knob, while Y453F is located in the RBD base and E484K and F486L are localized in the tip (**Fig. 2a**). F486 buries the greatest amount of surface area, of any residue, into the RBD-ACE2 complex. Although Y453, E484, and N501 are located in the ACE2 binding site, they form limited interactions with ACE2, defined by lower levels of buried surface area and only N501 forms a main chain hydrogen bond with ACE2 (**Fig. 2**).

Overall, the data show RBD residues that make limited contacts with ACE2 exhibit higher mutation frequencies than those involved in extensive hydrogen bond networks. For example, four SARS-CoV-2 S RBD residues (R403, S477, G485, and F490) were not counted in the ACE2 binding site because they bury less than 2Å^2^ of surface area into the complex. However, 3 of these residues, S477 (2.5%), G485 (0.01%), and F490 (0.018%), exhibit high mutation frequencies, showing that significant amino acid variation often “surrounds” residues critical for ACE2 binding.

### NAb/RBD structures for classification of NAb RBD binding epitopes

Several studies have classified SARS-CoV-2 RBD-binding NAbs (9,25), but an objective clustering rationale, requiring limited investigator interpretation, has not been described. As a result, a database of human NAb/RBD complexes (n=45) was compiled from the protein data bank (pdb) and used to characterize NAb RBD binding epitopes (**Table 1**). To define distinct features of each NAb/RBD complex, a binding profile (BP) was defined as the RBD residue-specific surface area buried upon NAb/RBD complex formation. NAb BPs were subjected to hierarchical clustering and compared using a similarity matrix (**Fig. 3**). The analysis provided an unbiased classification of anti-RBD NAbs into one of 5 distinct groups C1, C1D, C2, C3, and C4. Structures of all NAb/RBD complexes in each class are shown superimposed upon one another in **Figure. 4** and demonstrate striking structural conservation (e.g. C1) and diversity of NAb epitopes characterized by structural methods.

**Fig. 3.**
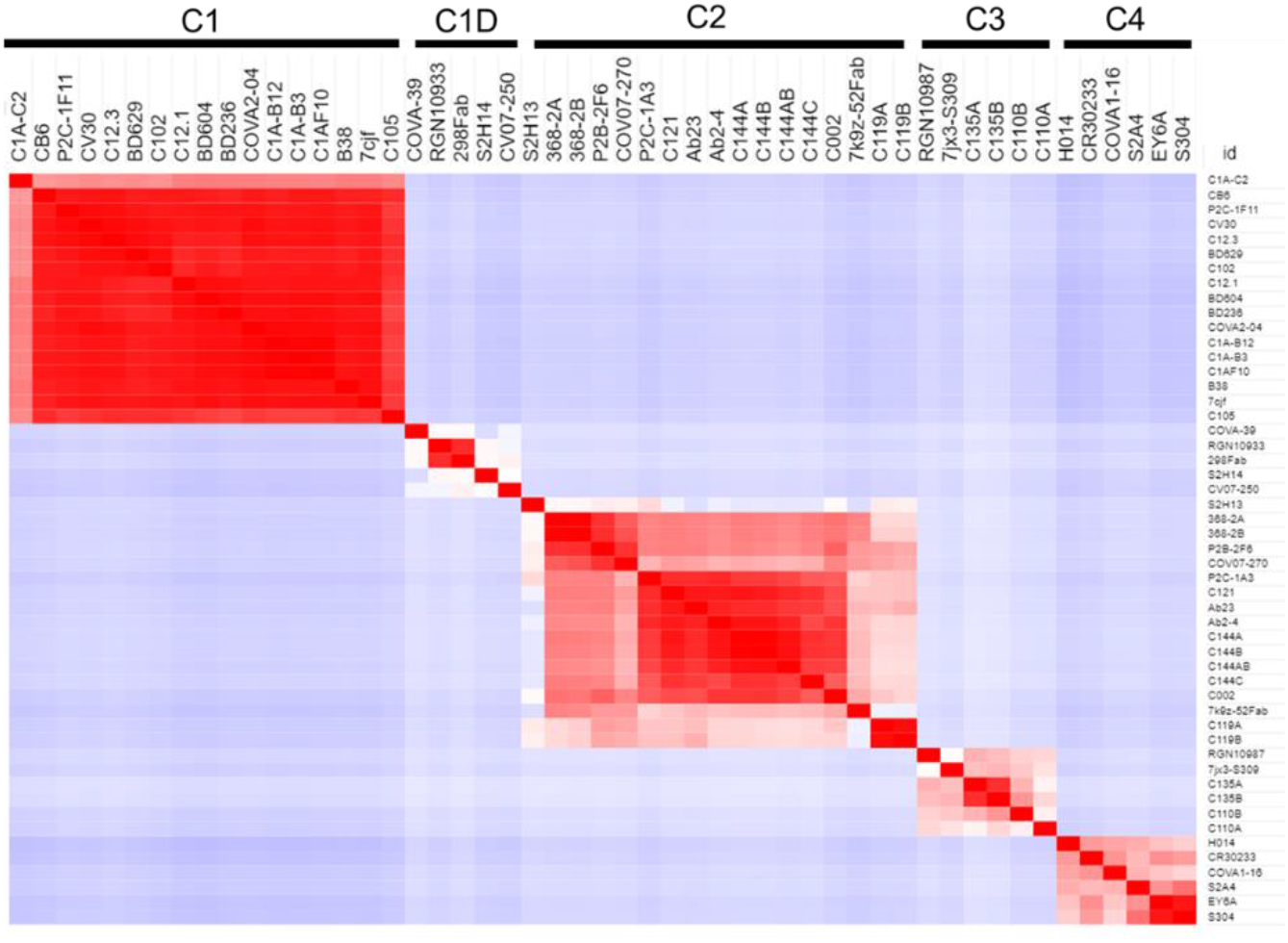
Binding Profile (BP) correlations to define RBD-binding NAb classes. BPs were defined for NAb/RBD complexes (Table 1) that were extracted from the pdb (n=45). For some structures, more than one NAb-RBD complex was evaluated leading to the A, B, C designations for some NAbs. Buried surface area was not calculated for some complexes, due to almost complete lack of sidechains for the NAbs. The BPs were subjected to hierarchical clustering and then similarity analysis (correlation coefficient calculation) to define five NAb classes C1 (n=17), C1D (n=5), C2 (n=12), C3 (n=4), and C4 (n=6).

**Fig. 4.**
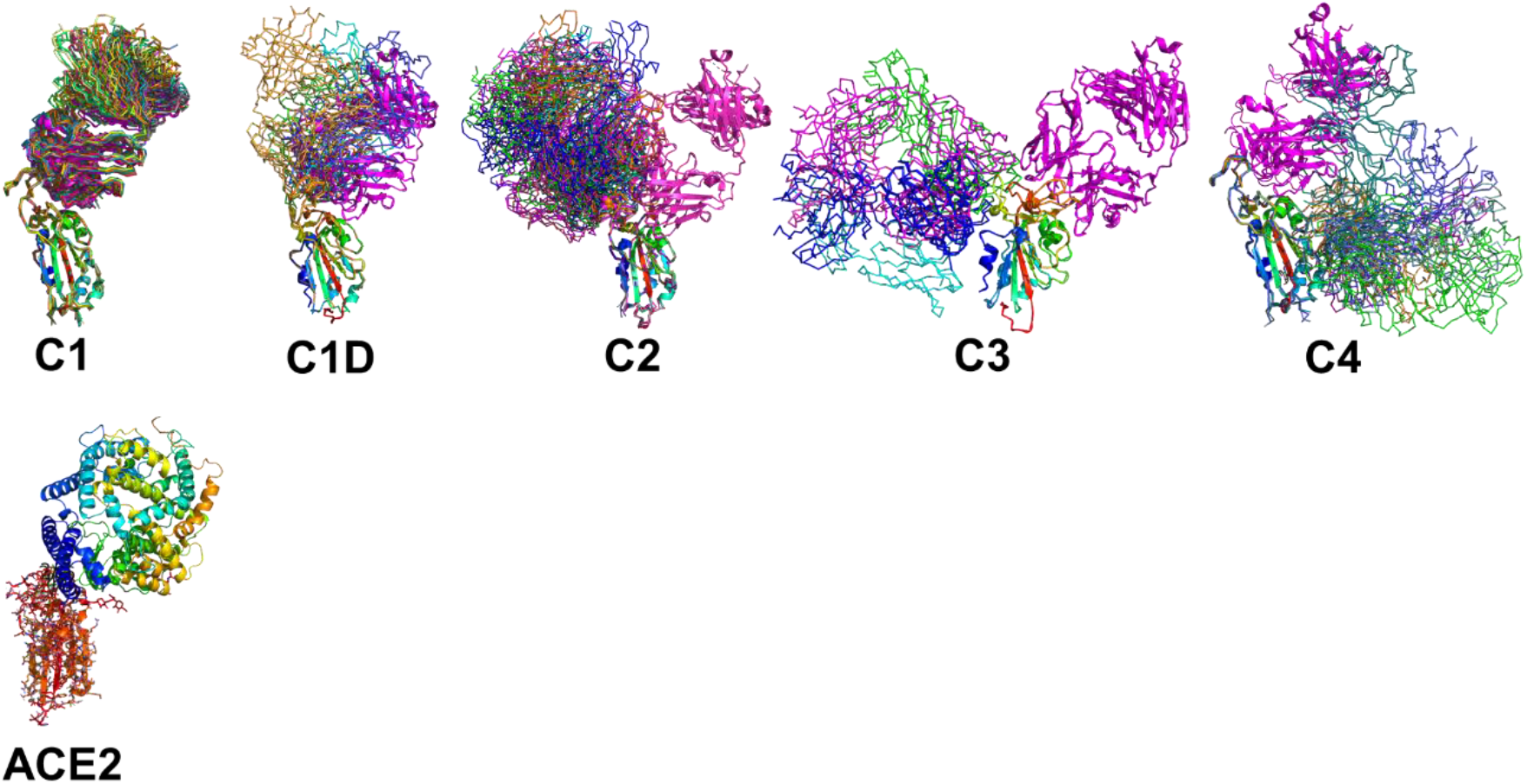
Epitope conservation and diversity within each NAb class. The RBD-ACE2 interaction, as well as all members of each NAb class are shown on the RBD. A ribbon diagram of an archetypical C1 NAb (magenta) is shown in each figure as a structural reference.

A total of 17 C1 NAbs were found in the pdb. C1 NAbs show striking structural conservation of their orientations and RBD-binding epitopes, which overlap substantially with the ACE2 binding site. As a result, C1 NAbs only bind to RBD when it is in the “up” conformation in the S protein (**Fig. 1**). BP analysis identified a distinct group of C1-like NAbs (C1D, n=5) that do not conserve the orientation or epitope of the C1 class (**Fig. 4**). C1D NAbs often bind closer to the flexible RBD tip (e.g. RGN10933 (4)) compared to C1 NAbs, which allows them to bind to RBD without it adopting the full “up” conformation required for C1 NAbs. Similarly, the steric disruption of C1D NAb COV7-250 binding to RBD in the “down” conformation is minimal (26). Thus, at least some C1D NAbs can bind to RBD with only minimal S conformational changes, yet their epitopes still extensively overlap with the ACE2 binding site.

Thirteen examples of C2 NAbs were identified, which show even greater diversity in their binding epitopes and orientations than C1 and C1D NAbs (**Fig. 4**). In fact, C1, C1D, and C2 NAb epitopes show a progression of binding epitopes and orientations that extend from one side of RBD that forms the ACE2 binding site (e.g. C1 NAbs), to the surface opposite the ACE2 binding site, where C2 NAbs bind without hindrance to the RBD in either up or down conformations. Thus, the accessibility of the C2 NAb epitopes are greater than for C1 and C1D NAbs. As previously reported, the orientations of several C2 NAbs, including Ab2-4, C121, and C144, shows they can block adjacent RBDs in the pre-fusion trimer from adopting the “up” ACE2-binding-competent conformations (9,27).

The C3 NAb (n=4) binding epitope is localized to the RBD knob and adjacent residues 333-347. C3 NAbs adopt distinct orientations, relative to the C1-C1D-C2 NAbs (**Fig. 4**). A unique feature of C3 NAb epitope is its location near RBD-RBD trimer interfaces. As a result, some C3 NAbs bind across adjacent RBDs in the trimeric SARS-CoV-2 S and sterically block the RBD up conformation of adjacent RBDs. Some C3 NAbs only block the up conformation of the RBD, while others sterically block ACE2 binding, and some have both neutralization mechanisms. Like C2 NAbs, C3 NAb epitopes are accessible whether SARS-CoV-2 S RBD is in the up or down conformation. C4 NAbs (n= 6) bind to an RBD surface comprising a helix-strand-helix motif (residues 366-389) that is buried in the trimeric S, even when the RBD is in the up conformation. Thus, C4 NAbs, required SARS-CoV-2 S to undergo significant conformational changes, beyond RBD up-down movements for binding (11). For many C4 NAbs, their epitope and orientation prevent them from sterically blocking ACE2 binding. This may explain why many C4 NAbs, such as CR3022, do not potently neutralize SARS-CoV-2 compared to NAbs in the other classes. However, this is not universal, as the orientations C4 NAbs COVA1-16 and H014 allow them to block ACE2 binding and they are reported to potently neutralize SARS-CoV-2 in cell culture assays and in animal models (28,29).

### Classification of NAbs isolated from a convalescent SARS-CoV-2 patient

We previously isolated a series of potent SARS-CoV-2 NAbs from a single convalescent patient and characterized their efficacy *in vitro,* and for one NAb (1212C2), its efficacy in golden Syrian hamsters, delivered by inhalation (16). The classified NAb/RBD structures (**Table 1, Fig. 4**) were used to predict the binding epitopes of these NAbs and the resulting assignments were experimentally tested using RBD variant binding and epitope mapping studies. Thus, unknown NAb heavy chains were aligned to the NAb database sequences classified as C1, C1D, C2, C3, or C4 (**Figs. 3 and 4**). Of 20 NAbs evaluated, 9 (45%) were classified as C1, 10 (50%) were classified as C2, and one (5%) was classified as C4. The analysis suggested that none of the unknown NAbs were C3 NAbs, and one of the 20 NAbs (1213H7) was misclassified by sequence analysis as a C2 NAb, but subsequently reassigned to the C1D classification based on epitope mapping and RBD variant binding analysis (**Figs. 6 and 7**). Alignments performed with the light chains did not improve the unknown NAb assignments.

### Susceptibility of NAbs to RBD variants

The predicted impact of RBD variant residues (**Fig. 2**) on NAb binding was evaluated using the BPs, derived from the NAb/RBD complex structures (**Table 1**). Variable RBD residues (freq. >0.01) found in at least 75% of the epitopes for each NAb class were identified (**Table 2**). This analysis identified 11 residues exhibiting increased variability in C1 and C1D NAb epitopes, 7 residues in the C2 epitope, and 2 in C3 NAb epitope. Variable residues that consistently bury surface area in at the C1 NAb class include K417, Y453, L455, A475, F486, and N501. Thus, C1 NAb epitopes are predicted to be sensitive to K417N/T, Y453F, L455F, A475V, F486L, and N501Y variants. Consistent with the structural relationships between C1, C1D, and C2 NAb epitopes (**Fig. 4**), conserved residues within the C1D epitope include residues shared with C1 NAbs (e.g. L455, A475, and F486), as well unique residues E484, and G485. Thus, RBD variants that could disrupt C1D NAbs include L455F, A475V, F486L, as well as E484K, and G485R. Contacts conserved across C2 NAbs include variable RBD residues G446, L452, V483, E484, F490, and S494. This is consistent with many C2 NAbs being sensitive to the E484K variant (9) and **Fig. 5**). Additional conserved RBD contacts within the C2 NAb epitope suggests RBD variants G446V, L452R, and S494P can also impact C2 interactions. Based on only four examples, the C3 NAb epitope is diverse and the conserved C3 contact region exhibits limited RBD residue variation. Only two RBD residues in the C3 epitope (N440 and K444) exhibit residue variability greater than 0.01%. Five additional RBD amino acids are contacted by 75% of the C3 NAbs, but these amino acid sidechains are highly conserved in the SARS-CoV-2 RBD sequences evaluated. In contrast, amino acid variation within the C1, C1D, and C2 epitopes is much greater (maximal variation 0.12%-2.5%) and extends over more residues. Finally, there are several common contact residues in the C4 NAb epitope, suggesting that N370S, T376I, V382L, P384L, T385I, and R408I could disrupt C4 NAbs.

**Table 2.** SARS-CoV-2 RBD residues conserved in ≥75% of NAb epitopes and the frequencies of variants.

|  | C1 | Avg. BSA | % Cons. | Variant | Freq. |
| --- | --- | --- | --- | --- | --- |
| 1 | Y505 | 97.2 | 100 | W | 0.004% |
| 2 | K417 | 80.2 | 100 | N | 0.024% |
| 3 | F486 | 61.6 | 100 | L | 0.059% |
| 4 | F456 | 57.1 | 100 | - | - |
| 5 | Y489 | 50.9 | 94 | - | - |
| 6 | R403 | 50.9 | 100 | I | 0.004% |
| 7 | A475 | 49.1 | 100 | S | 0.019% |
| 8 | Y421 | 48.0 | 100 | - | - |
| 9 | K458 | 45.1 | 100 | M | 0.001% |
| 10 | Q493 | 41.0 | 100 | K | 0.004% |
| 11 | L455 | 39.4 | 94 | F | 0.015% |
| 12 | N487 | 37.2 | 100 | H | 0.001% |
| 13 | T415 | 33.1 | 94 | M | 0.001% |
| 14 | Q498 | 31.6 | 88 | - | - |
| 15 | G502 | 28.6 | 100 | - | - |
| 16 | Y473 | 27.5 | 94 | - | - |
| 17 | Y453 | 27.3 | 100 | F | 0.062% |
| 18 | N460 | 26.3 | 100 | H | 0.002% |
| 19 | N501 | 25.0 | 100 | Y | 0.208% |
| 20 | G476 | 24.0 | 100 | S | 0.012% |
| 21 | G416 | 21.4 | 100 | - | - |
| 22 | S477 | 20.5 | 100 | N | 2.546% |
| 23 | T500 | 19.5 | 88 | - | - |
| 24 | D420 | 17.8 | 94 | - | - |
| 25 | D405 | 15.2 | 94 | - | - |
| 26 | Y495 | 13.1 | 88 | - | - |
| 27 | S459 | 11.2 | 100 | F | 0.021% |
| 28 | R457 | 10.7 | 100 | - | - |
| 29 | G496 | 10.2 | 76 | - | - |
| 30 | S494 | 8.7 | 88 | P | 0.046% |
| 31 | Q474 | 8.7 | 100 | - | - |
| 32 | F490 | 7.5 | 76 | S | 0.018% |

|  | C1D | Avg. BSA | % Cons. | Variant | Freq. |
| --- | --- | --- | --- | --- | --- |
| 1 | F486 | 152.3 | 100 | L | 0.059% |
| 2 | Y489 | 77.9 | 100 | - | - |
| 3 | Q493 | 60.7 | 100 | K | 0.004% |
| 4 | E484 | 54.1 | 100 | K | 0.079% |
| 5 | F456 | 48.7 | 100 | - | - |
| 6 | Y449 | 40.5 | 80 | - | - |
| 7 | N487 | 35.7 | 100 | H | 0.001% |
| 8 | L455 | 34.0 | 100 | F | 0.015% |
| 9 | S477 | 33.2 | 80 | N | 2.546% |
| 10 | A475 | 33.0 | 100 | V | 0.012% |
| 11 | G485 | 27.4 | 100 | R | 0.010% |
| 12 | Q498 | 26.7 | 80 | - | - |
| 13 | T478 | 19.1 | 80 | I | 0.063% |
| 14 | K417 | 17.8 | 80 | N | 0.024% |
| 15 | G476 | 14.7 | 80 | S | 0.012% |
| 16 | Y453 | 11.3 | 100 | F | 0.062% |
| 17 | S494 | 5.0 | 80 | P | 0.046% |

|  | C2 | Avg. BSA | % Cons. | Variant | Freq. |
| --- | --- | --- | --- | --- | --- |
| 1 | Y449 | 88.8 | 100 | - | - |
| 2 | E484 | 85.1 | 100 | K | 0.079% |
| 3 | F490 | 53.6 | 92 | S | 0.018% |
| 4 | Q493 | 39.1 | 92 | K | 0.004% |
| 5 | V483 | 29.0 | 92 | A | 0.020% |
| 6 | G446 | 24.9 | 92 | V | 0.027% |
| 7 | S494 | 23.1 | 92 | P | 0.046% |
| 8 | L452 | 22.5 | 92 | R | 0.121% |
| 9 | G485 | 20.4 | 83 | R | 0.010% |
| 10 | L492 | 8.0 | 100 | - | - |

|  | C3 | Avg. BSA | % Cons. | Variant | Freq. |
| --- | --- | --- | --- | --- | --- |
| 1 | R346 | 69.2 | 100 | K | 0.009% |
| 2 | T345 | 61.9 | 75 | - | - |
| 3 | L441 | 44.6 | 100 | - | - |
| 4 | N440 | 40.9 | 75 | K | 0.016% |
| 5 | K444 | 24.9 | 75 | R | 0.010% |
| 6 | N450 | 18.3 | 75 | - | - |
| 7 | N448 | 15.4 | 75 | - | - |

Red highlights frequencies $> 0.01\%$
Yellow corresponds to RBD variants evaluated for NAb and ACE2 binding

**Fig. 5.**
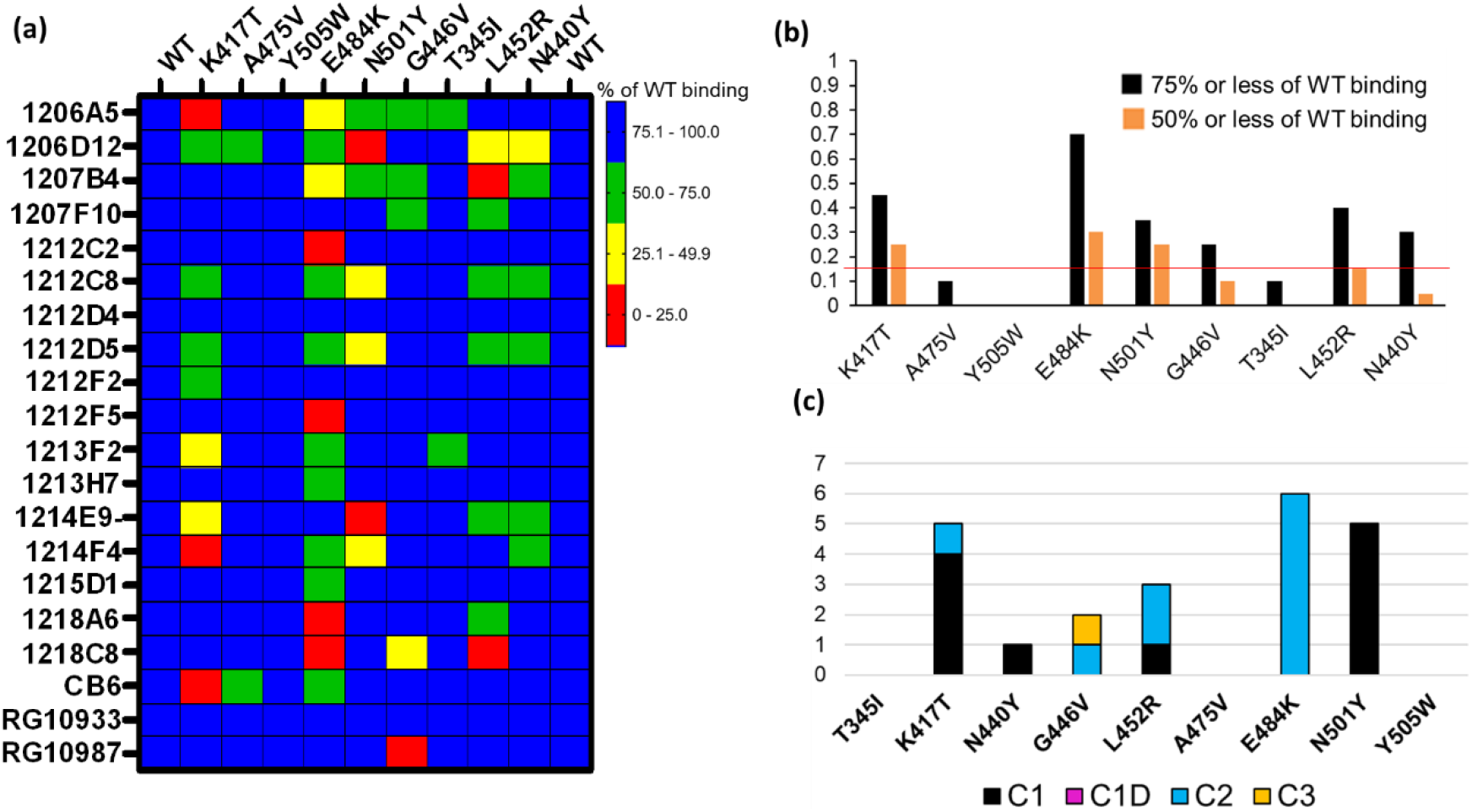
NAb binding to RBD variants. **(a)** SARS-CoV-2 RBD variants were injected over NAbs captured on a biacore chip and their binding levels measured and normalized to the binding response to a reference (WT) Wuhan-1 RBD. Two WT RBD binding experiments were performed at the beginning and end of the experiment to validate the reproducibility of the assay. NAb binding is presented in the matrix as % of WT binding. NAbs that exhibiti robust binding to all RBD variants are highlighted with red arrows. **(b)** Global summary of NAb binding to SARS-CoV-2 RBD variants. Black bars show the fraction of NAbs that exhibit red, yellow, or green levels of binding as shown in **Fig. 5a**. Orange bars represent the fraction of NAbs exhibit red and yellow binding, as shown in **Fig. 5a.** The horizontal red line highlights the RBD variants (K417T, E484K, and N501Y) that disrupt the binding of the largest number of NAbs. **(c)** Sensitivity of NAb classes to the different RBD variants. The number of NAbs that display less than 50% of WT binding in **Fig. 5a** (red and yellow boxes) were counted (y axis) and plotted according to their NAb classification (Table 1).

### RBD variant binding analysis

Based on the *in silico* NAb epitope analysis, nine RBD variants that localize to the C1 (K417T, A475V, N501Y, Y505W), C1D/ C2 (G446V, L452R and E484K) and C3 epitopes (T345I, N440Y, and G446V) were produced for binding studies with the unknown NAbs. C4 NAbs were excluded from the analysis. Our structural analysis, and recent literature (9), confirm that the loss of NAb binding to certain SARS-CoV-2 RBD variants can be used to identify NAb classes. Furthermore, the assays can be used to identify NAbs that are not sensitive to RBD variants and thus potentially useful as therapeutics. To address these questions, the unknown NAbs and three controls (CB6 (30), RGN10933 and RGN10987(4)) were tested for their ability to bind to the SARS-CoV-2 RBD variants (**Fig. 5**). NAb-RBD interactions were defined as “significantly disrupted” if NAb binding to an RBD variant was reduced by 50% or more, relative to the reference WT RBD. Based on this definition, C1 RBD variants K417T (25%) and N501Y (25%), and C2 variant E484K (30%), most frequently disrupted NAb binding (**Fig. 5**). A second group of RBD variants, L452R (15%), G446V (10%), and N440Y (5%) disrupted fewer NAb interactions, while all NAbs bound efficiently (>50%, relative to WT reference) to RBD variants T345I, A475V, and Y505W. Efficient binding of all NAbs to T345I (freq. = 0.0003%) and Y505W (freq. = 0.004%) is consistent with their high conservation within the SARS-CoV-2 RBD (**Table 2**). Furthermore, for those NAbs exhibiting >50% reduction in binding levels, relative to WT RBD, E484K and N501Y exclusively disrupted NAbs classified as C2 and C1, respectively (**Fig. 5c**).

### ACE2-RBD variant binding analysis

In addition to disrupting NAb binding, SARS-CoV-2 RBD mutations can impact RBD-ACE2 binding affinity, which could impact virus infectivity. Thus, ACE2 binding affinity was measured for each RBD variant and the reference Wuhan-1 RBD protein (**Fig. 6**). The two RBD variants that disrupted the largest number of NAb interactions (E484K, and N501Y) bound to ACE2 within 15% (e.g. error of the measurements) of the WT RBD-ACE2 interaction. In contrast, K417T, which disrupts a salt bridge with ACE2 (**Fig. 2b**), exhibited two-fold lower binding affinity for ACE2, relative to WT RBD (**Fig. 6b**). This data suggests 2 of the 3 most disruptive RBD mutations are NAb evading mutations that do not alter interactions with the viral receptor, ACE2. T345I and N440Y also exhibited WT RBD-ACE2 binding affinity, while G446V and A475V exhibited disrupted ACE2 binding affinities, like K417T. Two RBD mutations (L452R and Y505W) exhibited higher affinity for ACE2 than WT RBD. Thus, L452R disrupts several NAbs, but also binds to ACE2 with higher affinity. Thus, the L452R mutant, associated with a recently identified California VoC (B.1.427/B.1.429-California) has evolved to enhance ACE2 binding while simultaneously disrupting a significant number of NAbs in the C2 classification. Y505W also enhances RBD-ACE2 interactions but was not disruptive to any of the NAbs tested (**Fig. 5a**).

**Fig. 6.**
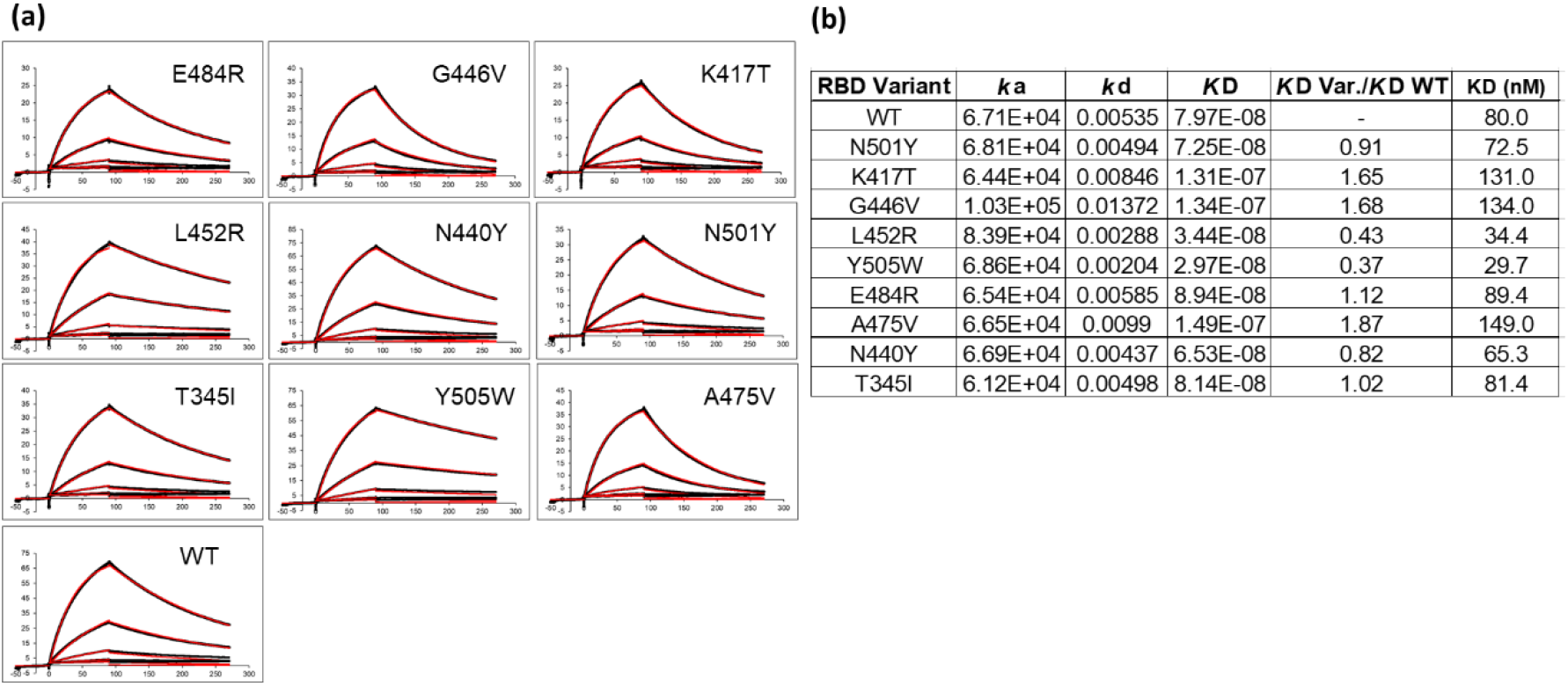
ACE2 binding to SARS-CoV-2 S RBD variants. **(a)** Sensorgrams of RBD variants binding to ACE2. **(b)** Binding constants derived from the sensorgrams.

### NAb Epitope Analysis

Epitope mapping was performed to further validate the assignment of the NAbs to the different classes. Based on their 50% neutralization (NT_50_) values and RBD variant binding properties, 12 NAbs were subjected to epitope mapping analysis (**Fig. 7**). Overall, there was excellent correlation between the NAb assignments (**Table 1**) and the epitope analysis. For example, C1 vs. C1 NAbs, or other “like-like” classifications prevented binding of the second NAb. In contrast, some C1 NAbs allowed simultaneous binding of a second C2 NAb, but this was not universal. For example, C1 NAbs 1212D5, 1212F2, 1213F2, 1212D4, and 1212C8, allowed simultaneous binding of C2 NAbs 1207B4, 1215D1, and 1207F10 (**Fig. 7**). In contrast, the C2 NAb 1212C2 blocked all C1 and C2 NAbs that were assayed (**Fig. 7a**). To address these observations, structures of the NAb-RBD complexes assigned to the different C2 NAbs were compared (**Fig. 7d**). Specifically, 1215D1, 1207B4, and 1207F10 NAbs were assigned to the C386-2/RBD complex, while 1212C2 was assigned to the C121/RBD complex. As shown in **Figure 7d**, 1212C2 (modeled by C121-RBD) overlaps with C1 (modeled by P2C-1F11/RBD) and C2 NAbs, while 1215D1 (modeled by C386-2/RBD) can simultaneously bind with several C1 (1214D4, 1212C8, 1212D5 etc.) and C1D (1213H7) NAbs. Thus, the assignment of each NAb to a NAb/RBD complex provided a structural basis for understanding the overlapping and distinct epitopes of the unknown NAbs

**Fig. 7.**
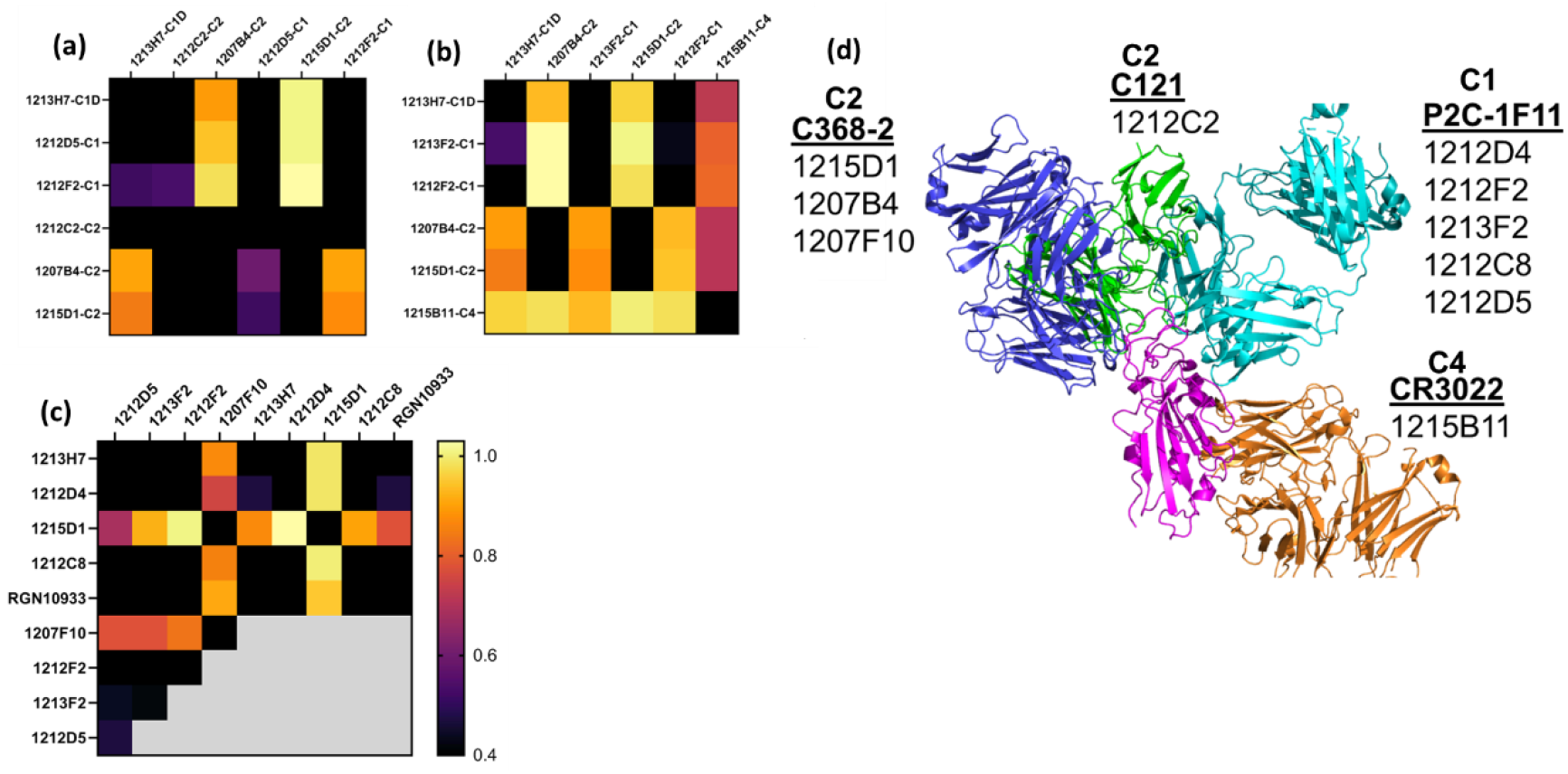
NAb epitope mapping. SPR-based epitope mapping performed to further validate the classification of unknown NAbs assigned to an NAb/RBD complex (Table 1). Figures **7a**, **7b**, and **7c**, represent 3 distinct mapping experiments performed with common and distinct NAbs. (**d**) Graphical summary, showing NAb/RBD complexes that were assigned to the unknown NAbs and used to explain the epitope mapping data. P2C-1F11/RBD was used to model C1 NAbs, C2 NAbs were modeled by C368-2/RBD (1215D1, 1207B4 and 1207F10) and C121/RBD (1212C2) structures, while the C4 NAb, 1215D11, was modeled by the RBD/CR3022 structure. The assigned structures explain how 1212C2 epitope blocks all other NAbs from binding to RBD, while 1212D4 (or 1213H7) and 1215D1 bind to non-overlapping RBD epitopes and also bind to all RBD variants they were tested against (**Fig. 5**). Two NAbs evaluated in the mapping data (1213H7 and ctl RGN10933) are not listed on the graphical summary. 1213H7 and RGN10933 were both assigned as C1D NAbs, which is consistent with their common epitope mapping profiles, and their ability to bind simultaneously with 1215D1.

As a second method of epitope analysis, the NAbs were tested for their ability to bind to an RBD-ACE2 fusion protein (FP). The RBD-ACE2FP was designed to provide a stable RBD-ACE2 complex to characterize the overlap of NAb epitopes with the RBD-ACE2 binding site. As expected, most NAbs bound very poorly to the RBD-ACE2FP, consistent with their epitope being blocked by ACE2 (**Fig. 8**). However, three C2-classified NAbs (1215D1, 1207B4, and 1207F10) exhibited high levels of binding to the RBD-ACE2FP (**Fig. 8**). Based on the location of the RBD ACE2 binding site (**Fig. 4**), this data further validates the assignment of these NAbs to the C2 class (**Fig. 7d**). It also suggests that this subclass of C2 NAbs may disrupt RBD-ACE2 interactions even after the complex is formed, presumably by a conformational change, providing a novel mechanism of neutralization for this group of C2 NAbs.

**Fig. 8.**
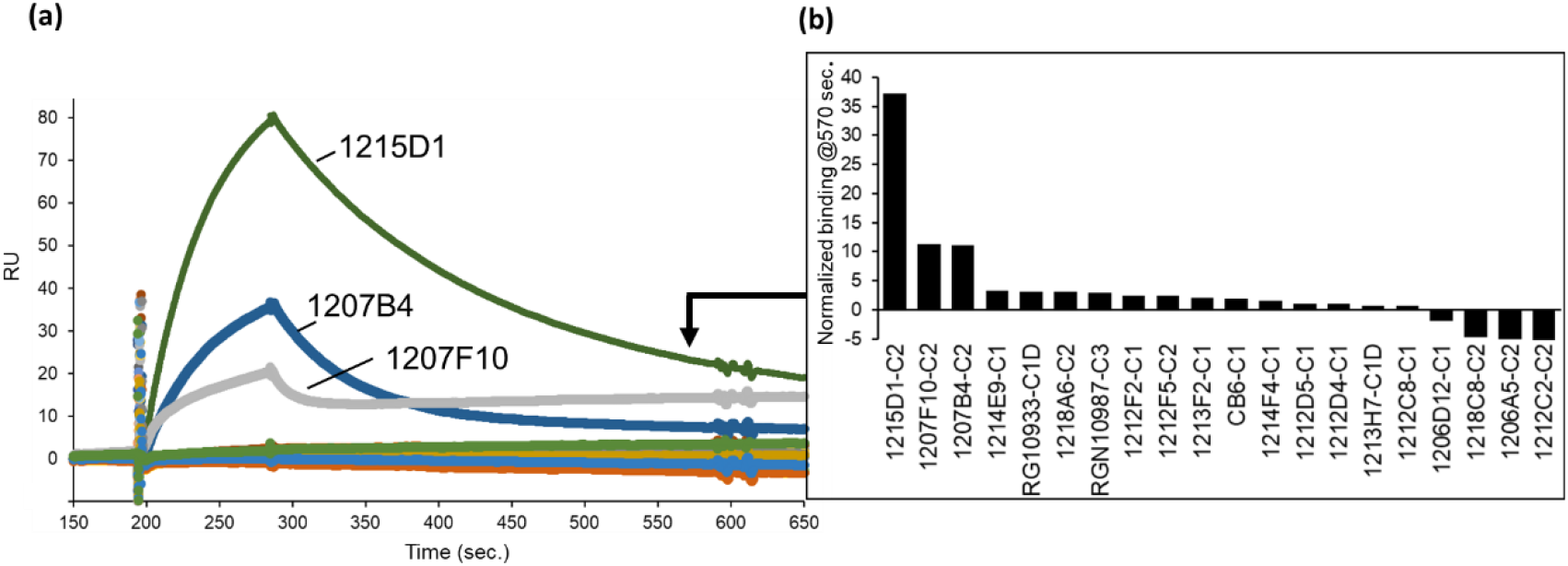
NAb epitope analysis using a SARS-CoV-2 RBD-ACE2 fusion protein. NAbs were captured on anti-IgG biacore chips and sensorgrams were recorded for an RBD-ACE2 fusion protein (20nM) injected over the surface of each NAb. Variability in fusion protein binding levels near the end of the sensorgrams (t = 570 seconds) is plotted in the graph.

## Discussion

The goal of this study was to use SARS-CoV-2 NAb/RBD structural information found in the protein databank and SARS-CoV-2 S RBD sequence variation data from the GSIAD, to characterize a series of NAbs for potential use as therapeutics against SARS-CoV-2. Optimal NAbs against SARS-CoV-2 have always required that they exhibit high potency neutralizing capabilities, but as the pandemic has continued, they now must also exhibit broad specificity against naturally occurring SARS-CoV-2 VoC. The current VoC include B.1.1.7-UK (N501Y, S494P*, E484K*), P.1-Japan/Brazil (K417N/T, E484K, N501Y), B.1.351-South Africa (K417N, E484K, N501Y) B.1.427/B.1.429-California (L452R), where amino acid changes in the SARS-CoV-2 S RBD of these VoC are listed in parentheses (cdc.gov).

RBD amino acid mutations found in the VoC, with the exception of the K417T mutation (K417T frequency = 0 in the analysis on 1-11-21), are found in the top 20 of most frequently observed mutations (frequency > 0.024%) in the GSIAD. SARS-CoV-2 VoC have been shown to reduce Ab-mediated neutralization and/or exhibit increased disease severity and transmissibility (cdc.gov). To further evaluate these single mutations at the molecular level, we tested a panel of SARS-CoV-2 NAbs, isolated from a convalescent patient, for their ability to bind to a series of SARS-CoV-2 RBD variants. The analysis found that E484K is the most disruptive mutation and impacted the greatest number of NAbs tested in our study (30%), with N501Y and K417T being slightly less disruptive, but still each impacting 25% of the NAbs evaluated. Combined, the 3 SARS-CoV-2 RBD variants (K417T, E484K, and N501Y) present in the P.1 Japan/Brazil and B.1.351 South Africa VoC significantly disrupted the binding of 65% of the NAbs tested.

In addition to disrupting NAb binding, which is correlated with reduced virus neutralization, the impact of the variants on ACE2 binding affinity was determined. This is an important question, often not addressed, that may define mechanisms of viral selection responsible for increased infectivity and/or disease severity, as well as disrupting host NAb responses. Analysis of RBD-variant/ACE2 binding affinity revealed that N501Y and E484K exhibited essentially WT affinity for ACE2, where WT corresponds to the Wuhan-1 SARS-CoV-2 RBD sequence. RBD variants N440Y and I345I also exhibited ACE2 binding affinity similar to WT RBD. This suggests that neither N501Y and/or E484K variants increase infectivity, reported for the B.1.1.7 UK VoC, by increasing affinity for ACE2. If increased ACE2 affinity is required for increasing viral infectivity, other mutations in the SARS-CoV-2 S, such as D614G, P681H, or several other changes, alone or together, must be responsible for enhanced ACE2 binding affinity. Overall, our data suggests that N501Y and E484K enhance viral fitness predominantly by disrupting the host NAb response, but not by an increase in ACE2 binding affinity.

Like N501Y and E484K, K417T also significantly disrupts NAb binding. However, the K417T variant, as well as the G446V and A475V variants exhibit reduced (~2-fold) ACE2 binding affinity. Thus, escape from NAb binding/neutralization at these positions comes with a cost of reducing ACE2 binding affinity, suggesting these mutants are unlikely to confer a selective advantage to the virus though enhanced viral attachment. In contrast, L452R and exhibited a 2-fold increase in ACE2 affinity. This is consistent with the B.1.429 California VoC being able to evade NAb responses, as well as enhance ACE2 binding affinity, possibly providing advantages in immune evasion against NAbs, and increased cell infectivity via enhanced ACE2 binding affinity, ultimately leading to increased disease severity. A weakness in our studies is that our RBD-ACE2 binding analysis studies are based on a 1:1 monomeric ACE2-RBD interaction. It is possible that the conformation and function of the RBD variants are distinct in the context of the trimeric SARS-CoV-2 S. Recent data suggests that the D614G mutation stabilizes the S trimer and is necessary to observe the functional impact of the N501Y and L452R mutations on cell infectivity (31,32). It is also possible that large changes in RBD-ACE2 affinity, higher or lower, disrupts optimal viral infectivity, while more subtle changes around the reference WT RBD-ACE2 binding affinity can cause large changes in cell infectivity.

From an overall analysis of SARS-CoV-2 anti-RBD NAbs, C3 NAbs appear to have an advantage by targeting an RBD epitope that has undergone less variation than NAb epitopes associated with C1, C1D, and C2 NAbs. However, variability in CDR length, structure, and amino acid composition has thus far been sufficient for at least some C1, C1D, and C2 NAbs to maintain their neutralization efficacy, despite RBD variability. Furthermore, we find the C3 RGN10987 NAb is highly sensitive to the G446V variant, suggesting the virus is able to identify residues to disrupt NAbs in all NAb classes. For the unknown NAbs evaluated in this report, the NAb/RBD database, RBD variant binding studies, and epitope mapping allowed their assignment into specific NAb classes. Thus, our study provides a strategy for characterizing NAb/RBD epitopes at a larger scale to further define NAb sensitivity to RBD variants in individuals recovered from natural infections and changes in the anti-RBD SARS-CoV-2 structural repertoire induced by vaccination. Finally, our analysis identified a non-overlapping antibody pair, 1213H7/1215D1, which both bind to all RBD variants found in current SARS-CoV-2 VoC, suggesting they represent a potential therapeutic treatment for SARS-CoV-2 WT and newly identified VoC.

## Ethics statement

A.D., J.J.K., L.M.S., J.J.K, and M.R.W. are co-inventors on a patent for NAbs described in the study.

## Acknowledgements

We acknowledge use of the UAB Heflin Genomics Core, UAB Multidisciplinary Molecular Interaction Core Facility.

## Author Contributions

M.R.W. and A.D. conceived the study, A.D., B.D.H., and M.R.W. performed experiments, A.D., M.R.W., L.M.S., and J.J.K. wrote and edited the paper.

